# Syngeneic model of carcinogen-induced tumor mimics basal/squamous, stromal-rich, and neuroendocrine molecular and immunological features of muscle-invasive bladder cancer

**DOI:** 10.1101/2022.12.03.518865

**Authors:** Shruti D Shah, Bryan M Gillard, Michelle M Wrobel, Ellen Karasik, Michael T Moser, Michalis Mastri, Norbert Sule, Craig M Brackett, Wendy J Huss, Barbara A Foster

## Abstract

**Background:** Bladder cancer is a heterogenous disease and the emerging knowledge on molecular classification of bladder tumors could have impact to drive treatment decisions based on molecular subtype. Pre-clinical models representing each subtype are needed to test novel therapies. Carcinogen-induced bladder cancer models represent heterogeneous, immune-competent, pre-clinical testing options with many features found in the human disease.

**Methods:** Invasive bladder tumors were induced in C57BL/6 mice when continuously exposed to N-butyl-N-(4-hydroxbutyl nitrosamine) (BBN) in the drinking water. Tumors were excised and serially passed by subcutaneous implantation into sex-matched syngeneic C57BL/6 hosts. Eight tumor lines were developed and named BBN-induced Urothelium Roswell Park (BURP) tumor lines. The BURP lines were characterized by applying consensus molecular classification to RNA expression, histopathology, and immune profiles by CIBERSORT. Two lines were further characterized for cisplatin response.

**Results:** Eight BURP tumor lines were established with 3 male and 3 female BURP tumor lines, having the basal/squamous (BaSq) molecular phenotype and morphology. BURP-16SR was established from a male mouse and has a stromal-rich (SR) molecular phenotype and a sarcomatoid carcinoma morphology. BURP-19NE was established from a male mouse and has a neuroendocrine (NE)-like molecular phenotype and poorly differentiated morphology. The established BURP tumor lines have unique immune profiles with fewer immune infiltrates compared to their originating BBN-induced tumors. The immune profiles of the BURP tumor lines capture some of the features observed in the molecular classifications of human bladder cancer. BURP-16SR growth was inhibited by cisplatin treatment, while BURP-24BaSq did not respond to cisplatin.

**Conclusions:** The BURP tumor lines represent several molecular classifications, including basal/squamous, stroma-rich, and NE-like. The stroma-rich (BURP-16SR) and NE-like (BURP-19NE) represent unique immunocompetent models that can be used to test novel treatments in these less common bladder cancer subtypes. Six basal/squamous tumor lines were established from both male and female mice. Overall, the BURP tumor lines have less heterogeneity than the carcinogen-induced tumors and can be used to evaluate treatment response without the confounding mixed response often observed in heterogeneous tumors. Additionally, basal/squamous tumor lines were established and maintained in both male and female mice, thereby allowing these tumor lines to be used to compare differential treatment responses between sexes.

## Introduction

Bladder cancer is histologically and molecularly a heterogenous disease. Tumor molecular heterogeneity has been evaluated by genomic and transcriptomic analysis which led to the development of a “consensus” molecular classification system (1). The human bladder cancer “consensus” molecular classification system provides six clusters that take into account various different transcriptomic classification which may be used for treatment selection and improved therapeutic outcomes (1, 2). The six molecular subtypes in the consensus molecular classification, arranged from most to least differentiated, are Luminal Papillary (LumP) 24%, Luminal Non-Specified (LumNS) 8%, Luminal Unstable (LumU) 15%, Stroma-rich (SR) 15%, Basal/Squamous (Ba/Sq) 35%, and Neuroendocrine-like (NE-like) 3% (1). Patients show differences in survival hazard ratios based on their molecular subtype. The median overall survival is highest for the more differentiated LumP at 4 years and SR at 3.8 years. In contrast, the median overall survival for patients with the less differentiated Ba/Sq and NE-like tumors is only 1.2 and 1 year, respectively (1). We have previously used the “consensus” molecular classification in patient-derived xenograft (PDX) bladder cancer models to determine how representative each model is of the human disease. The human patient-derived xenograft models were classified as Ba/Sq or LumP even though some tumors demonstrated epithelial mesenchyme transition (EMT) and NE pathologic features (3). PDX models associated with different molecular subtypes are important tools for evaluating treatment response and novel targeted therapeutics that allow personalized treatment options for patients based on their molecular subtype.

In addition to patient-derived tumor models, carcinogen-induced models are useful, especially in organ sites with higher risks associated with carcinogen exposure. The N-butyl-N-(4-hydroxybutyl)-nitrosamine (BBN) carcinogen-induced mouse bladder tumor model is well established and forms invasive bladder tumors with mutation patterns that closely resemble the mutation pattern of human bladder cancer (4). BBN is a nitrosamine alkylating compound closely related to smoking carcinogens and specifically induces urothelial cancers in rodents when added to the drinking water (5). Analysis of BBN carcinogen-induced bladder tumors indicates that these tumors are transcriptionally and histologically similar to human tumors (4, 6). Muscle invasive tumors develop after approximately 12-20 weeks of exposure to the carcinogen (4, 7, 8). BBN-induced tumors show characteristic changes of non-muscle invasive (NMIBC) and muscle-invasive bladder cancer (MIBC). The BBN carcinogen-induced model of bladder cancer has been used in preclinical prevention, NMIBC, and MIBC studies. BBN initiates formation of bladder cancer, and the cancers progress even after BBN exposure stops (9, 10),(11). BBN-induced tumorigenesis is specific to the bladder. As seen in human bladder cancer the BBN-induced model has a sex disparity where male mice develop more tumors at shorter BBN exposure compared to female mice (12). Unfortunately, most recent studies in the BBN model only utilize the male mice. The mutation profile in early- and late-stage tumors was studied in male mice (4). Specifically, mutations in *Trp53* (80%), *Kmt2d* (70%), *Kmt2c* (90%), *Hmcn1* (90%), and *Arid1a* (30%) are frequently mutated in the BBN-induced tumors in male C57BL/6 mice (4). A comparison between BBN-induced bladder cancer in C57BL/6 and FVB host demonstrated that urothelial cell carcinoma often had squamous features in both strain backgrounds, but glandular differentiation was only found in the FVB strain (13). The inflammatory response in BBN-induced tumors was characterized by increased immunoinhibitory molecules leading to tumor escape (11).

One of the limiting factors to testing novel therapeutics in bladder cancer, including those that have an immune component to their efficacy such as immune checkpoint inhibitors, is the lack of preclinical models that reflect human disease. Optimally preclinical models should represent the molecular subtypes seen in patients, consider the male-to-female bladder cancer incidence ratio of 3:1, and reflect that the composition of immune cells infiltrating the tumor (immune contexture). The molecular subtype and immune contexture strongly associate with overall survival in bladder cancer (14); thereby highlighting the need for pre-clinical models reflecting the spectrum of clinical disease to develop novel therapeutics. The BBN carcinogen-induced model recapitulates many of these key features of the clinical disease including presenting with a variety of molecular subtypes, sex disparity with male mice presenting with disease early and female with more aggressive disease later and presenting with a variety of immune infiltrates. Unfortunately, the high degree of heterogeneity and long extended timeframe required for the development of tumors in the BBN carcinogen-induced mouse bladder tumor model presents many logistical challenges for *in vivo* mechanistic studies of novel therapeutics. In the current study, eight sex-matched BBN-induced Urothelium Roswell Park (BURP) tumor lines were developed from C57BL/6NTac male and female mice after continuous exposure to BBN in the drinking water. Because these models represent molecularly distinct subtypes of bladder cancer in both male and female mice, they provide unique models to study the impact of molecular subtype, sex disparities, and immune contexture on novel immune and non-immune based therapeutics in bladder cancer.

## Methods

### Summary of experimental design is depicted in Figure 1

#### Development of a syngeneic immune intact model of bladder cancer

##### BBN model and BBN-induced Urothelial carcinoma Roswell Park (BURP) tumor lines

Mice were housed in the Laboratory Animal Shared Resource (LASR) at Roswell Park in a limited access barrier facility. LASR is an AAALAC International Accredited Animal Program. Male and female C57BL/6NTac mice at 8 weeks of age received 0.1% of BBN (TCI Chemicals, Cat # B0938) in their drinking water *ad libitum* for up to 36 weeks. Weekly abdominal palpitations monitored bladder tumor formation. The high-grade disease was determined by detection of tumor on abdominal palpation and/or by detection of blood in the urine, at which time the mice were euthanized as per Roswell Park Institutional Animal Care and Use Committee (IACUC) guidelines. Bladders were removed and processed for passage subcutaneously into sex-matched C57BL/6NTac host mice. Mice treated with BBN that developed primary bladder tumors were denoted as donor mice. The tumors that grew from the first subcutaneous passage from donor mice were referred to as P0 passage, and the subsequent passages as P1, P2, etc. The allograft models were considered stable BURP tumor lines after no histological drift was observed between passages, usually after P2.

**Figure 1.**
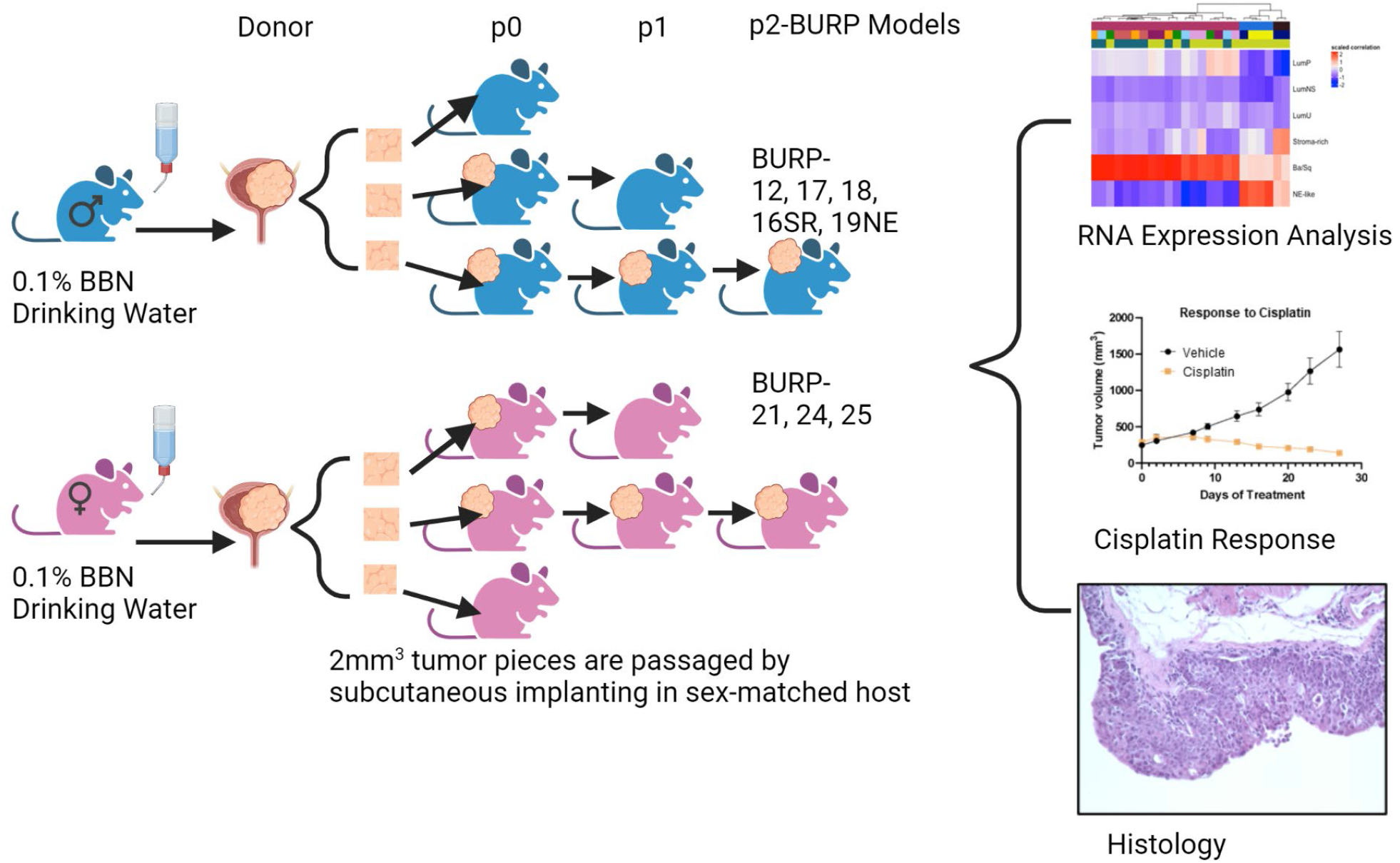
Graphical representation of methods. Created in BioRender.

##### Histologic evaluation and immunohistochemistry (IHC) analysis

Tissues were fixed in 10% buffered formalin for 24 hours before processing for paraffin embedding. Once fixed, samples were embedded in paraffin and sectioned at 5 microns on Starfrost adhesive slides (Mercedes Medical; Catalog #MER 7255/90/WH). Slides were deparaffinized in three xylene baths and then rehydrated in graded 100% to 70% alcohols, followed by ddH_2_O. IHC was performed using the DAKO Autostainer Plus, and Hematoxylin and Eosin (H&E) (Agilent Technologies, #CS11830-2) staining was performed using the DAKO CoverStainer. For IHC staining, slides were incubated in 1x pH6 citrate buffer (Invitrogen Cat #00-5000) for 20 minutes. Slides were incubated in 3% H_2_O_2_ (ThermoFisher Scientific; Catalog #H325-500) for 15 min to quench endogenous peroxidase activity. To block non-specific binding, tissues were incubated with normal goat or rabbit serum for 10 min (Table 1), followed by avidin/biotin block (Vector Labs Cat#SP-2001). Primary antibodies (Table 1) were diluted in 1% bovine serum albumin (BSA) solution (ThermoFisher Scientific; Catalog# BP1605-100) and incubated for 30 minutes at room temperature, followed by the biotinylated secondary antibodies (Table 1) for 15 minutes at room temperature. ABC reagent (Vector Labs Cat #PK 6100) was applied for signal enhancement for 30 minutes at room temperature. To reveal the peroxidase activity, slides were incubated with 3,3’-Diaminobenzidine (DAB) substrate (Dako Cat #K3467) for 5 minutes and counterstained with DAKO Hematoxylin for 20 seconds at room temperature. Slides for H&E and IHC were dehydrated through several baths of graded alcohols and three xylenes and then coverslipped using the DAKO CoverStainer.

**Table 1.**
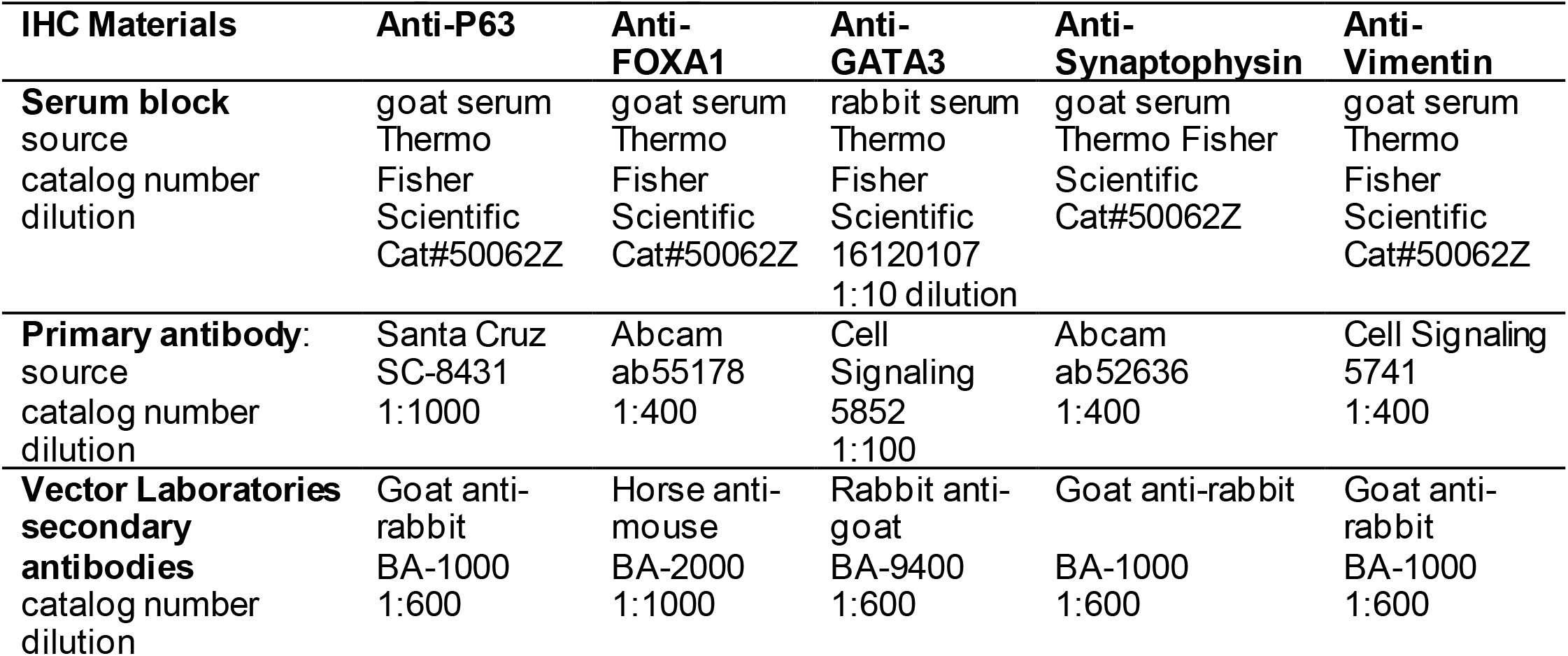
Immunohistological staining reagents, sources and dilutions.

#### Cisplatin treatment

BURP-16SR and BURP-24BaSq lines were implanted subcutaneously as 1-2 mm_3_ pieces into sex-matched C57BL/6NTac host mice (BURP-16SR n=21; BURP-24BaSq n =19). When tumors reached 250mm_3_, approximately 12 weeks after implantation, body weight was recorded weekly to monitor mice randomized into treated and control groups. Mice were treated with 10mg/kg of cisplatin 1 mg/ml formulation (TEVA Pharmaceuticals South Wales PA) or saline vehicle control, I.V. once a week for 4 weeks, and caliper measurements measured change in tumor growth. One week after final treatment, mice were collected for tumor analysis (BURP-16SR n=10 Cisplatin; n=11 Vehicle; BURP-24BaSq n =7 Cisplatin; n=8 Vehicle). Tumor growth curves of treated compared to vehicle controls were analyzed with unpaired student t-test using Prism GraphPad software.

#### Whole transcriptome sequencing (RNA-seq)

##### Total RNA isolation

Total and small RNA was isolated and purified using the miRNeasy mini kit (Qiagen, Cat # 217084) according to the manufacturer’s directions. Briefly, 10-50 mg of frozen tissue was homogenized for 5 minutes in 700 μl of Qiazol reagent using Navy Rhino tubes in a Bullet Blender Homogenizer (Next Advance). The homogenates were allowed to sit at room temperature for 5 mins. After incubation, chloroform was added, and the samples were shaken for 15 sec. After adding chloroform, the homogenates were separated into aqueous and organic phases by centrifugation. RNA partitions to the upper aqueous phase, DNA partitions to the interphase and proteins to the lower organic phase or the interphase. The upper aqueous phase was transferred to a fresh tube, and ethanol was added to provide appropriate binding conditions for all RNA molecules larger than 18 nucleotides. The aqueous fraction samples were then applied to the miRNeasy Mini spin column, where total RNA bound to the membrane and phenol and other contaminants were efficiently washed away. On-column DNAse digestion was performed to remove residual genomic DNA contamination, followed by additional washes. High-quality RNA was eluted in 60 μl of RNase-free water. A quantitative assessment of the purified total RNA was performed using a Qubit Broad Range RNA kit (Thermofisher Cat# Q10210). The concentration was determined by Ribogreen fluorescent binding to isolated RNA. The RNA was further evaluated qualitatively using RNA Nanotape on the 4200 Tapestation (Agilent technologies), where the sizing of the RNA was determined, and a qualitative numerical score (RINe) was assigned to each sample. RNA with RIN of greater than 7 was used for RNA-seq analysis.

##### Whole transcriptome sequencing (RNA-seq)

RNA-seq analysis was performed by the Roswell Park Genomics Shared Resource. The sequencing libraries were prepared with the RNA HyperPrep Kit with RiboErase (HMR) (Roche Sequencing Solutions) using 500 ng of total RNA, following the manufacturer’s instructions. Briefly, the first step depletes rRNA from total RNA, followed by DNA digestion to remove any gDNA contamination. Next, samples were purified, fragmented, and primed for cDNA synthesis. The fragmented RNA was reverse transcribed into first-strand cDNA using random primers. The next step removed the RNA template and synthesized a replacement strand, incorporating dUTP in place of dTTP to generate ds cDNA. Pure Beads (Kapa Biosystems) were used to separate the ds cDNA from the second strand reaction mix resulting in blunt-ended cDNA. A single ‘A’ nucleotide was added to the 3’ ends of the blunt fragments. Multiple indexing adapters, containing a single ‘T’ nucleotide on the 3’ end of the adapter, were ligated to the ends of the ds cDNA, preparing them for hybridization onto a flow cell. Adapter ligated libraries were amplified by PCR, purified using Pure Beads, and validated for appropriate size on a 4200 TapeStation D1000 Screentape (Agilent Technologies, Inc.). The DNA libraries were quantified using a Kapa Biosystems qPCR kit and pooled at an equimolar concentration. Each pool was denatured and diluted to 350 pM with a 1% PhiX control library added. The resulting pool was loaded into the appropriate NovaSeq Reagent cartridge for 100-cycle paired-end sequencing and run on a NovaSeq6000 following the manufacturer’s recommended protocol (Illumina Inc.). Sequencing quality control was assessed using FASTQC v0.11.5 (http://www.bioinformatics.babraham.ac.uk/projects/fastqc/). Reads were aligned to the mouse genome GRCM38 M16 (genocode) using STAR v2.6.0a (PMID 23104886) and post-alignment quality control was assessed using RSeQC v2.6.5 (PMID 22743226). Aligned reads were quantified using RSEM v1.3.1 (PMID 21816040).

#### Molecular analysis

##### Differential gene expression analysis

Raw reads from RNA-seq data were transformed to CPM and log-CPM by applying the CPM transformation function of the *edgeR* R package (Bioconductor). After the transformation, low expressed genes were filtered out and normalized by the trimmed mean of M-values. Multidimensional scaling (MDS) was performed using the *limma* R package (Bioconductor) to identify tumor lines that are transcriptionally similar and hence clustered together. The expression values of tumor lines were compared to normal mouse bladder sequence obtained from GSE112973 (7) by fitting linear models using the limma package to identify differentially expressed genes for each tumor line. The false discovery rate was less than 0.05. The regulon activity of 23 regulators associated with bladder cancer was determined by the following method as outlined by Aurelie and colleagues using the *RTN* R package (2.6.0) (1). Succinctly, a regulatory network is provided as an RTN TNI-class object, and the regulon activity is calculated for each tumor line. Using the RTN’s *tni*.*gsea2* function, the two-tailed GSEA tests were calculated. The enrichment score (dES) is the difference between the positive and negative enrichment score and represents regulon activity. Unsupervised hierarchical clustering was performed for the signature using the *ComplexHeatmap* R package.

##### Consensus molecular classification of BURP tumor lines

The molecular subtype of BURP tumor lines was determined by applying the consensus classifiers developed by Kamoun and colleagues, using the *consensusClassifier* R package (1). This package implements the nearest-centroid transcriptomic classifier. Since the classifier labels were specific for human Entrez IDs, the mouse gene ids were converted to homologous human gene ids using the *BiomaRt* R package. Using log-CPM values, the correlation values to the classifiers were obtained using a minimum correlation of 0.1.

##### Identifying immune cell profile of BURP tumor lines from bulk tumor expression data

The CIBERSORT algorithm (15) was applied to deconvolve the immune cell signature from bulk tumor expression data. For identifying the mouse immune cells, the input signature matrix from ImmuCC (16, 17) was used that included signatures of 25 murine immune cells. The CIBERSORT algorithm deconvolves the solid tumor by applying a machine learning approach called support vector regression method that solves for identifying the relative fraction of each cell type.

## Results

### BBN-induced Urothelial carcinoma Roswell Park (BURP) tumor model

The BURP tumor lines were developed by exposing C57BL/6NTac mice to 0.1% BBN carcinogen in the drinking water ad libitum until palpable tumors formed. C57BL/6NTac mice on 0.1% BBN drinking water developed spontaneous primary tumors in the bladder as early as 20 weeks, with high-grade muscle-invasive disease developing between 28 and 36 weeks, as determined by pathologic analysis. Stable tumor lines were developed by passaging primary carcinogen-induced tumors subcutaneously into sex-matched C57BL/6NTac immune-competent mice as 1-2 mm_3_ tumor tissue. A tumor line was considered stable when there was no drift in the histology from the previous passage number. In most cases, each BURP tumor line was considered stable after three passages (p2) or less. Eight tumor lines, 5 from male, and 3 from female, were established, and pathology was characterized in comparison to the original donor tumors (Table 2 and Figure 2). Histologically, all of the original BBN-induced bladder tumors were muscle-invasive with conventional urothelial carcinoma characteristics containing squamous differentiation and sarcomatoid features (Table 2 and Figure 2). All of the BURP tumor lines showed reduced differentiation and immune infiltrates compared to the original tumors as indicated by lymphocyte detection by pathologic analysis. Six of the eight BURP lines (BURP-12, -17, -18, -21, -24, -25) have similar pathology of urothelial carcinoma (with squamous differentiation) comparable to their originating tumors; while BURP-16SR has a spindling pleomorphic pathology and BURP-19NE demonstrated poorly differentiated pathology.

**Table 2.** Donor and BURP model sex, pathology, tumor stage, and immune infiltrates.

| **Line** | **Sex** | **Donor** |  |  | **BURP Line** |  |
| --- | --- | --- | --- | --- | --- | --- |
|  |  | **Pathology** | **Stage** | **Immune Infiltrates** | **Pathology** | **Immune Infiltrates** |
| **BURP-12** | M | Invasive UC, w/sq diff | T3 | Minimal | UC w/sq diff | Minimal |
| **BURP-16SR** | M | Invasive UC, with sarcomatoid features | T2 | Moderate | Sarcomatoid | Minimal |
| **BURP-17** | M | Invasive UC, w/sq diff | T2 | Minimal | UC w/sq diff | Minimal |
| **BURP-18** | M | Invasive UC, w/sq diff | T2 | Moderate | UC w/sq diff | Minimal |
| **BURP-19NE** | M | Invasive UC, w/sq diff | T1 | Heavy | Poorly diff | Minimal |
| **BURP-21** | F | Invasive UC, w/sq diff | T1 | Minimal | UC w/sq diff | Minimal |
| **BURP-24** | F | Invasive UC, w/sq diff and sarcomatoid features | T2 | Minimal | UC w/sq diff | Minimal |
| **BURP-25** | F | Invasive UC, w/sq diff | T1 | Minimal | UC with sq diff | Minimal |

Key: M=Male; F=Female; UC=urothelial cell carcinoma; w/ = with, sq diff = squamous differentiation

**Figure 2:**
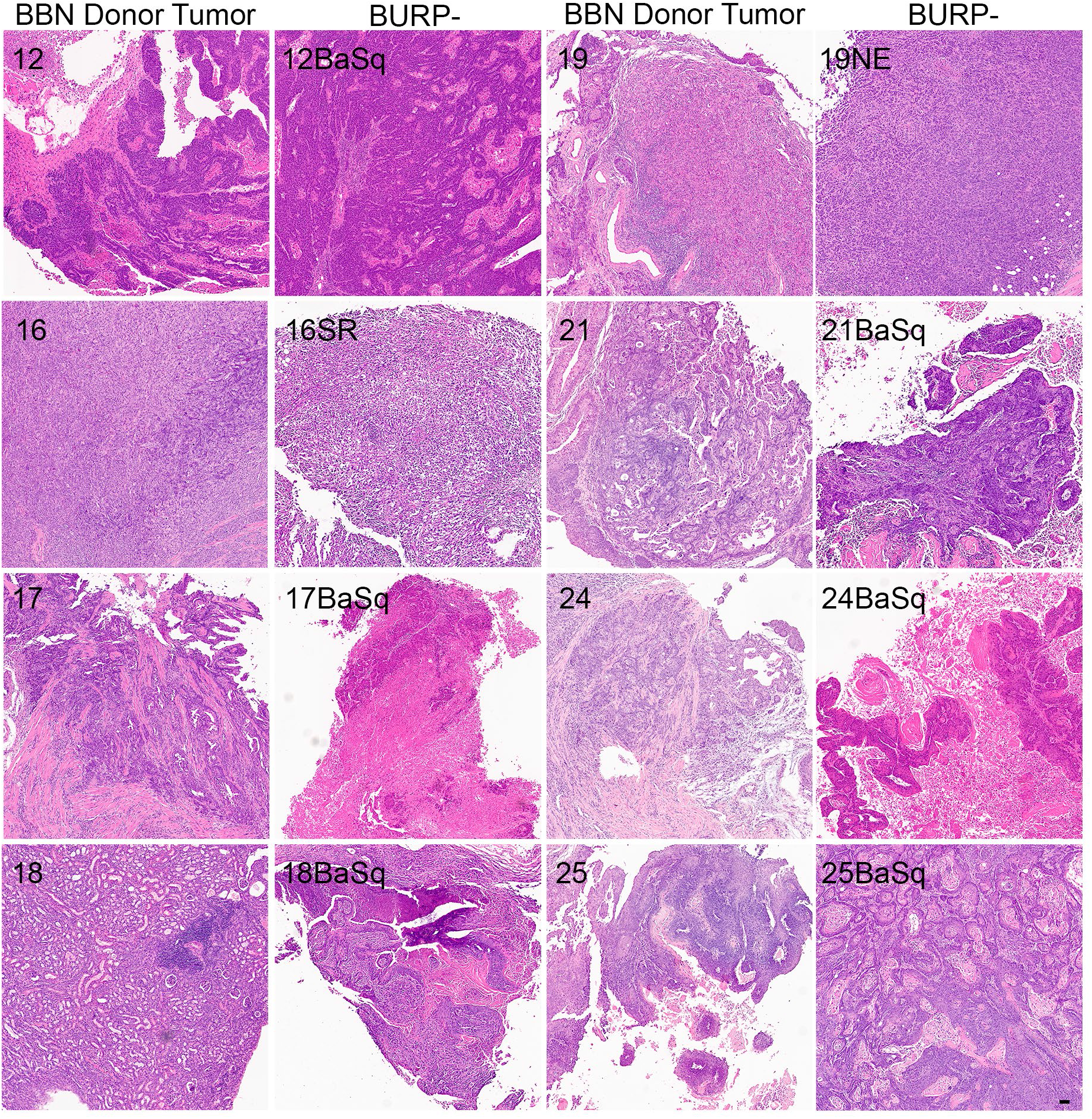
Generation of histologically diverse BURP tumor lines. H&E of original high-grade muscle-invasive tumors (BBN Donor Tumor) in male and female C57BL/6 mice exposed to 0.1% BBN. Representative H&E from each established BURP tumor lines from each original BBN Donor Tumor, 50μm scale bar is in the bottom right panel.

### Molecular features of BURP tumor models

Differential gene expressionof the BURP tumor lines was evaluated using transcriptomic data obtained via RNA-seq analysis of three independent tumors from each line at passages between p2 and p5, the time at which the tumor lines were considered stable. Multidimensional scaling (MDS) analysis was performed using bulk tumor RNA-seq data to identify the transcriptomic differences between the eight BURP tumor lines. The transcriptomic differences between the tumor lines were mapped using euclidean distance and represented as an MDS plot (Figure 3A). Each data point represents an individual tumor, with the color representing the BURP tumor line and the shape representing the sex of mouse from which the tumor line was derived. The MDS plot shows that the BURP tumor lines clustered in 3 groups, indicating that the eight BURP models can be divided into 3 transcriptomic classifications (Figure 3A). Among the three groups, Clusters 2 and 3 each had one BURP tumor line (BURP-19NE and BURP-16SR, respectively). Cluster 1 contains all six tumor lines with squamous differentiation (BURP-12BaSq, -17, -18, -21, -24, and -25).

**Figure 3:**
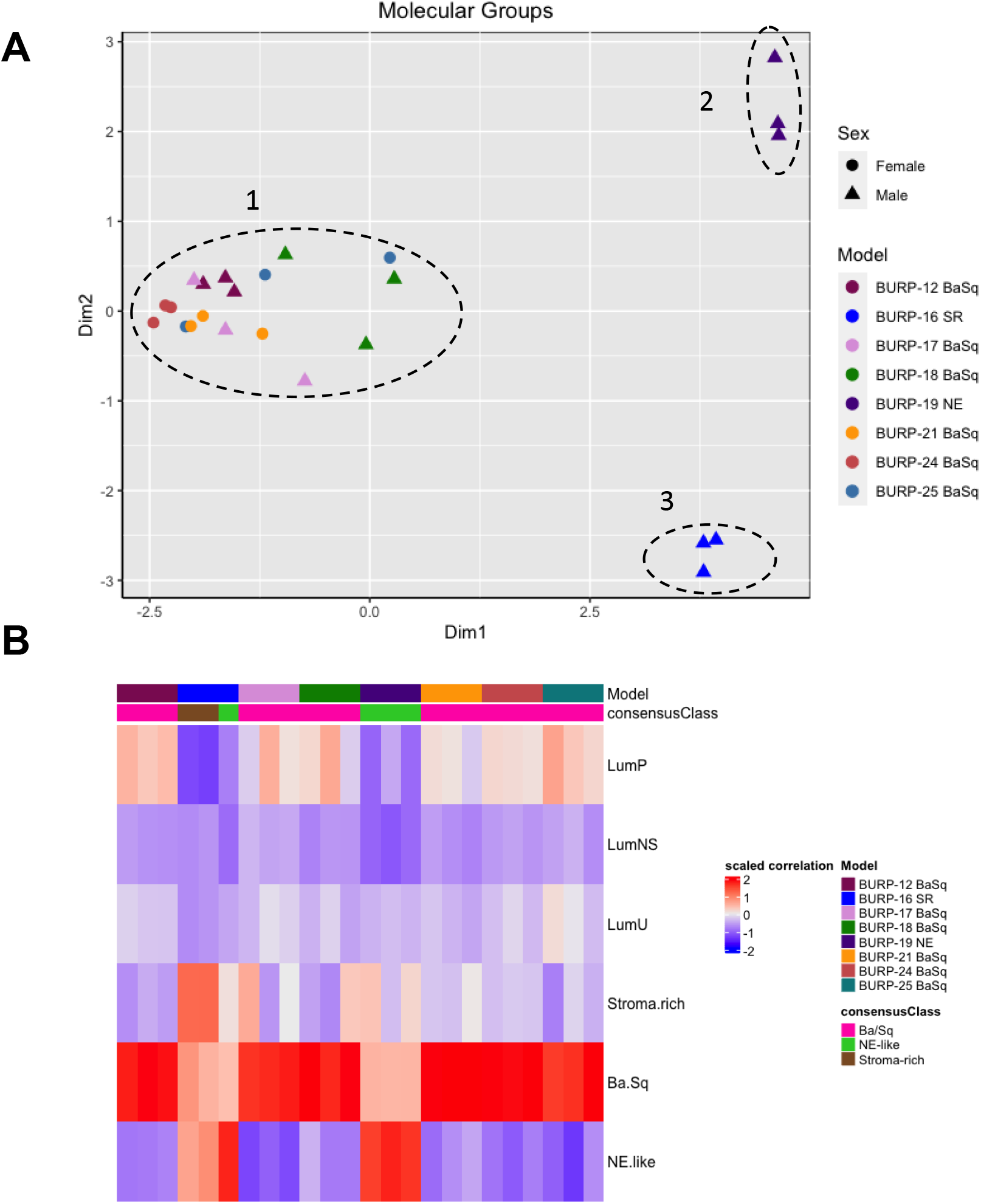
Molecular characterization of BURP tumor lines. **(A)** Multidimensional scaling (MDS) analysis of gene expression in the eight BURP tumor lines. Each data point represents an individual tumor with different colors representing different BURP tumor lines. Male lines are indicated by a triangle and female lines by a circle. The dotted lines mark three different clusters. The x and y axis depict eigenvalues in 2 dimensions (Dim1 and Dim2). **(B)** Heatmap of Molecular Consensus classifiers from Kamoun et al., 2020 for the BURP tumor lines. The color represents scaled correlation coefficient values between the BURP tumor line transcriptome data and the Consensus classifiers. Red represents higher correlation, and blue represents lower correlation.

Next the association of the BURP lines with the consensus molecular classifiers from Kamoun et al. (1) was determined. The consensus molecular analysis correlates the input expression matrix and the different molecular classifiers. The highest positive correlation between the input gene matrix and the molecular classifier helps identify the molecular classification for that particular tumor line. Of the six different molecular classifications, BURP tumor lines had a high positive correlation with the Ba/Sq, stroma-rich, and NE-like subtypes (Figure 3B). Six tumor lines (BURP-12, -17, -18, -21, -24, -25BaSq) of Cluster 1 had high correlation with Ba/Sq classification (Figure 3A) and had squamous differentiation morphology (Figure 2). BURP-19NE had high correlations with the NE-like classification (Cluster 2, Figure 3A) and a spindling pleomorphic pathology (Figure 2). BURP-16SR had a high correlation with stroma-rich (2/3 replicates) and NE-like (1/3 replicates) classifications (Cluster 3, Figure 3A) and poorly differentiated pathology (Figure 2). None of the tumor lines showed a high positive correlation with LumP, LumNS, or LumU classification.

### Differentiation marker expression in BURP tumor models

IHC of differentiation markers was used to confirm similarities of the BURP tumor lines analyzed with multidimensional scaling analysis and consensus molecular classification. IHC was performed to further characterize the BURP models for several markers of differentiation, including TP63 (basal cell) marker, FOXA1 (luminal cell), and GATA3 (luminal cell) (18, 19). The six Ba/Sq molecular subtype/Cluster 1 models expressed different patterns of TP63, FOXA1, and GATA3 (Figure 4). BURP-12 expressed TP63, FOXA1, and GATA3; BURP-17 and BURP-24 expressed FOXA1 and GATA3; BURP-18 and BURP-21 only expressed TP63; and BURP-25 expressed TP63 and FOXA1. To further characterize the BURP-16 and BURP-19 models, synaptophysin (NE-like) and Vimentin (EMT/stroma) markers were analyzed in addition to TP63, FOXA1 and GATA3 (Figure 5). In BURP-16 expression of FOXA1, synaptophysin and vimentin were high; TP63 was expressed by some cells; and GATA3 was not detected. In BURP-19 expression of synaptophysin was high, confirming the NE-like molecular subtype; TP63 and FOXA1 were also highly expressed, whereas GATA3 and Vimentin expression was more heterogeneous. Thus, the molecular and histological characterization of BURP tumor lines reveals three different subtypes of bladder cancer: Ba/Sq (BURP-12, -17, -18, -24, and -25BaSq); NE-like (BURP-19NE); and Stroma-rich/NE-like (BURP-16SR).

**Figure 4:**
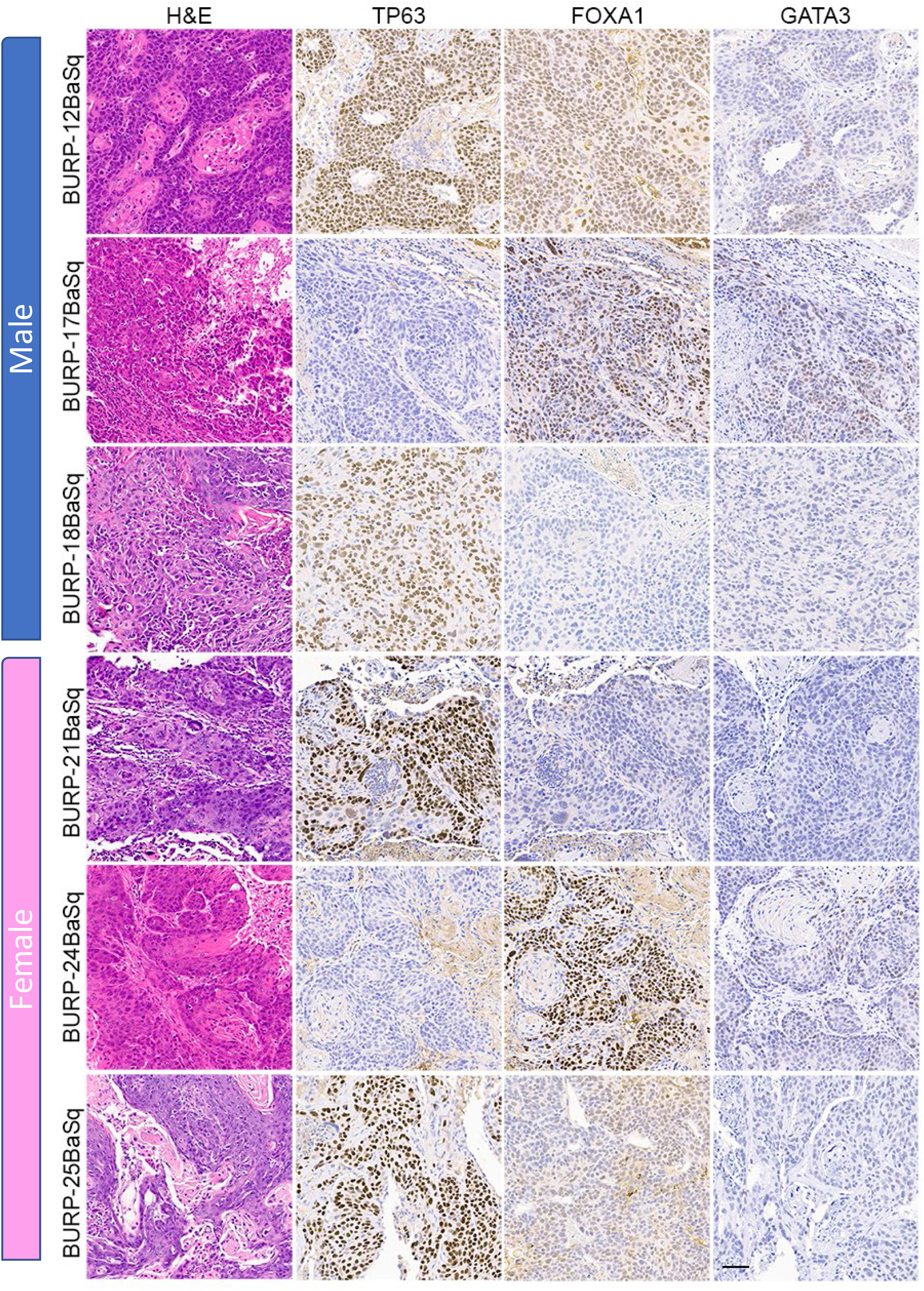
IHC of differentiation markers in the BURP lines with Ba/Sq molecular subtype. Serial sections of BURP-12, -17, -18, -21, -24, -25 were stained with H&E, TP63 IHC, FOXA1 IHC, and GATA3 IHC. Blue bar indicates male models and pink bar indicates female models. 50μm scale bar is in the bottom right panel.

**Figure 5:**
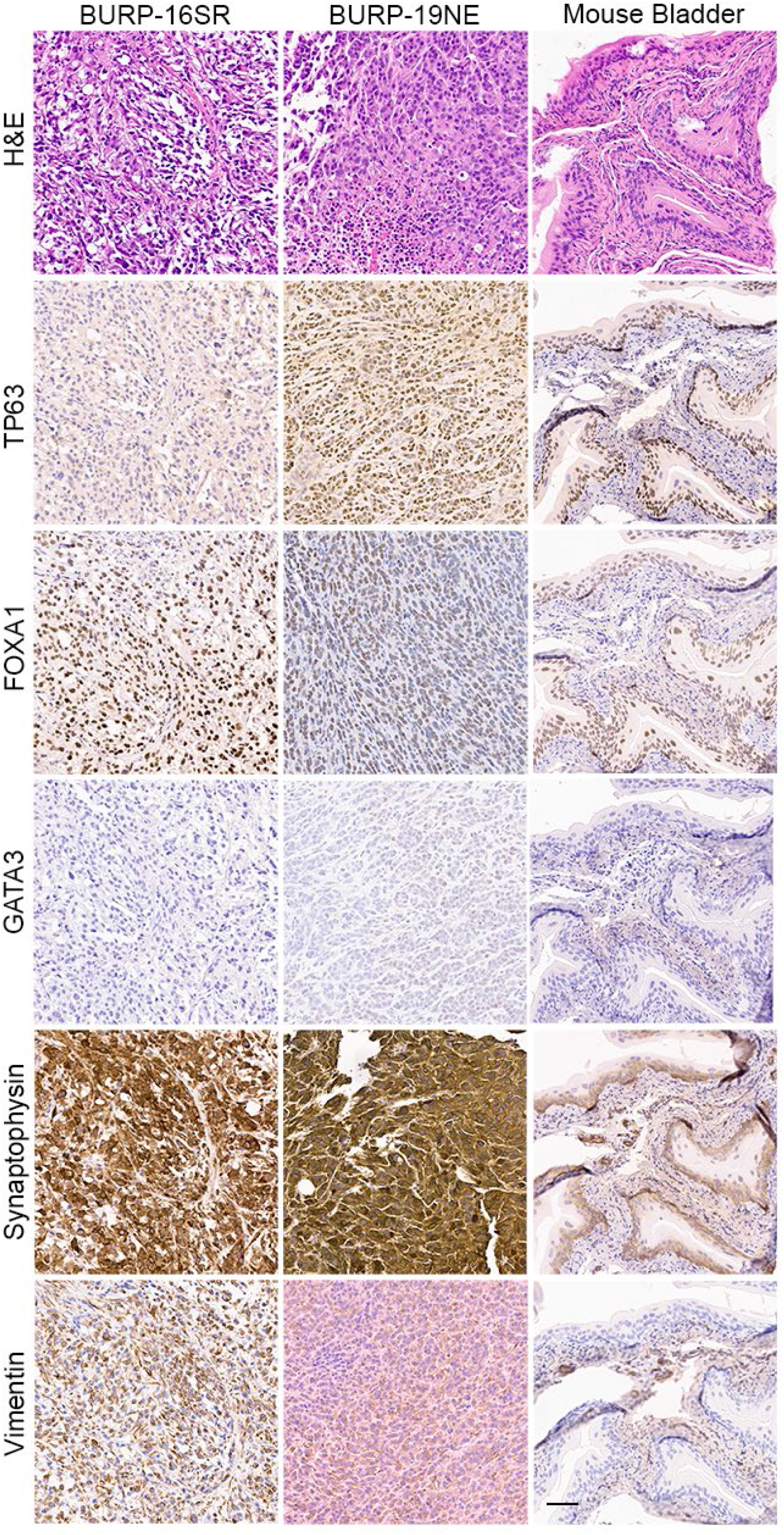
IHC of differentiation markers in the BURP-16SR and BURP-19NE lines. Serial sections were stained with H&E, TP63 IHC, FOXA1 IHC, GATA3 IHC, Synaptophysin IHC, and Vimentin IHC. Normal bladder control from BURP-16SR allograft is a control for the IHC staining. 50μm scale bar is in the bottom right panel.

### Immune contexture of BURP tumor lines

Molecular subtype specific differences in the immune contexture, which refers to the composition of the immune landscape in the tumor, is a predictor of survival in bladder cancer patients (20-23). For example, M1 polarized macrophages, activated dendritic cells (DCs), CD8^+^ tumor infiltrating lymphocytes (TILs), and anti-tumor helper CD4^+^ T cells (Th1) positively correlate with survival. In contrast, the presence of pro-tumorigenic M2 polarized macrophages and the CD4^+^ T cell helper subpopulations, T regulatory cells (Tregs), Th2, and Th17 cells associate with poor survival outcome. To understand whether molecular subtype was associated with a distinct immune contexture in the BURP lines, the deconvolution algorithm CIBERSORT using the ImmuCC immune cell signature was applied to bulk RNA-seq data that included 25 immune cell signatures (Figure 6). Macrophages, B cells, DCs, and T cells had relatively higher fractions across all BURP tumor lines.

**Figure 6:**
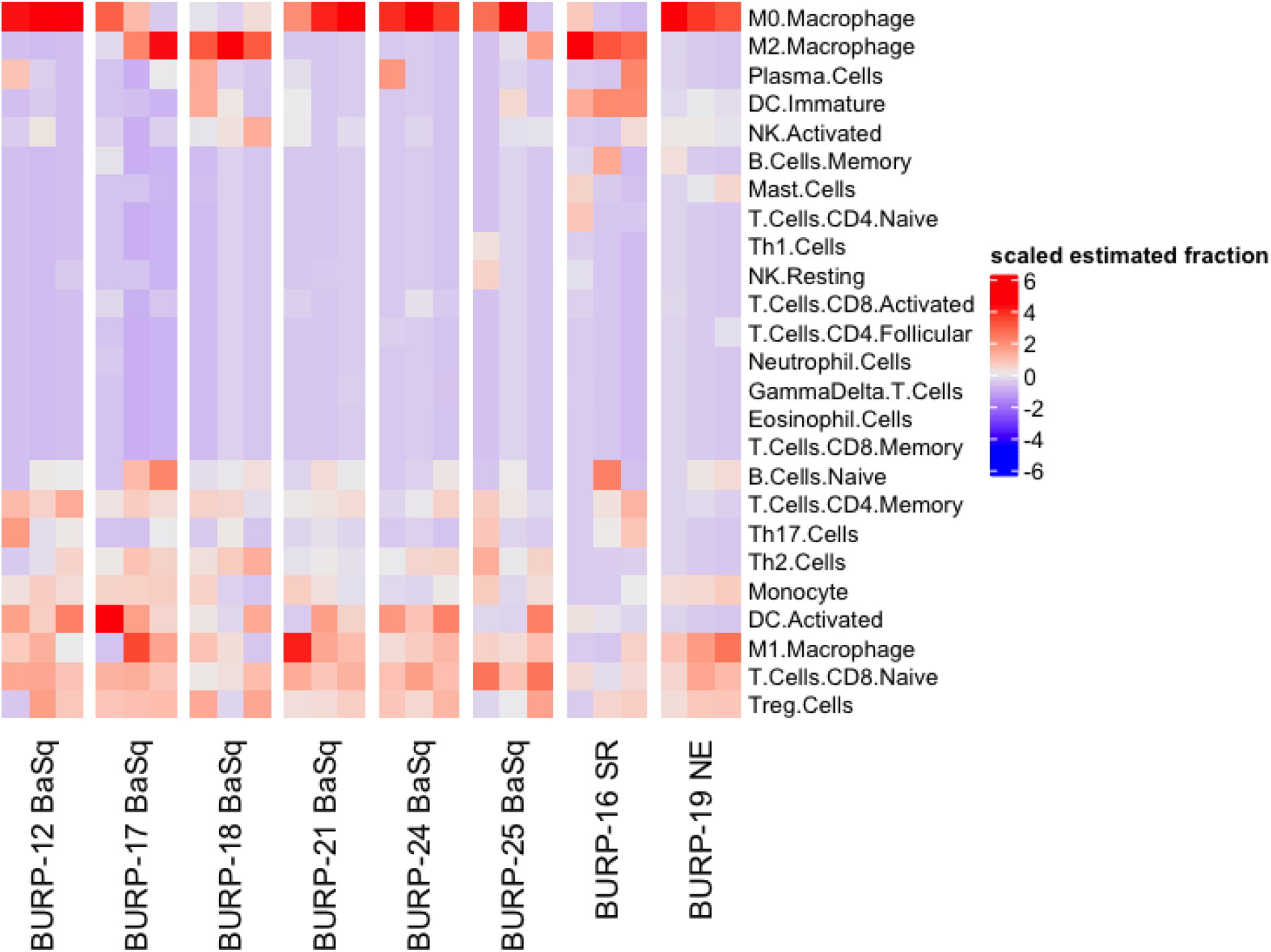
Immune contexture of BURP models. Spearman correlation analysis of ImmuCC cell signature of BURP tumor model. Red represents positive correlation and blue represents negative correlation.

The estimated fraction of macrophages, B cells, DCs, and T cells and the fraction of these immune populations with an activated status was examined more closely (Figure 7). Macrophages made up the highest estimated fraction among all the immune cell populations in all the BURP tumor lines. The fraction of non-polarized M0, anti-tumor M1, and pro-tumor M2 macrophages within the total macrophages was estimated for each BURP line (Figure 7A). In the Ba/Sq BURP lines, these macrophage populations fell into three categories: 1) higher M0 than M1 macrophages in BURP-12, -21, -24; 2) a mix of M0, M1, and M2 macrophages in BURP-17 and -25; and 3) predominately M2 macrophages in BURP-18. BURP-16SR consisted predominately of M2 macrophages. BURP-19NE consisted mostly of M0 macrophages with some M1 macrophages. When comparing the fractions of naïve and memory B cells (Figure 7B), the Ba/Sq BURP tumor lines had a higher fraction of naïve B cells, the BURP-16SR line had a mix of naïve and memory B cells, and the BURP-19NE line also had a mix of naïve and memory B cells. The estimated fraction of immature and activated DCs in the Ba/Sq BURP lines fell into two categories: 1) almost exclusively activated DCs in BURP-12, -17, -24; and 2) mix of immature and activated DCs in BURP-18, -21, and -25 (Figure 7C). BURP-16SR was made up of mostly immature DCs. BURP-19NE also consisted of immature DCs but had the smallest estimated fraction of total DCs of all BURP lines.

**Figure 7:**
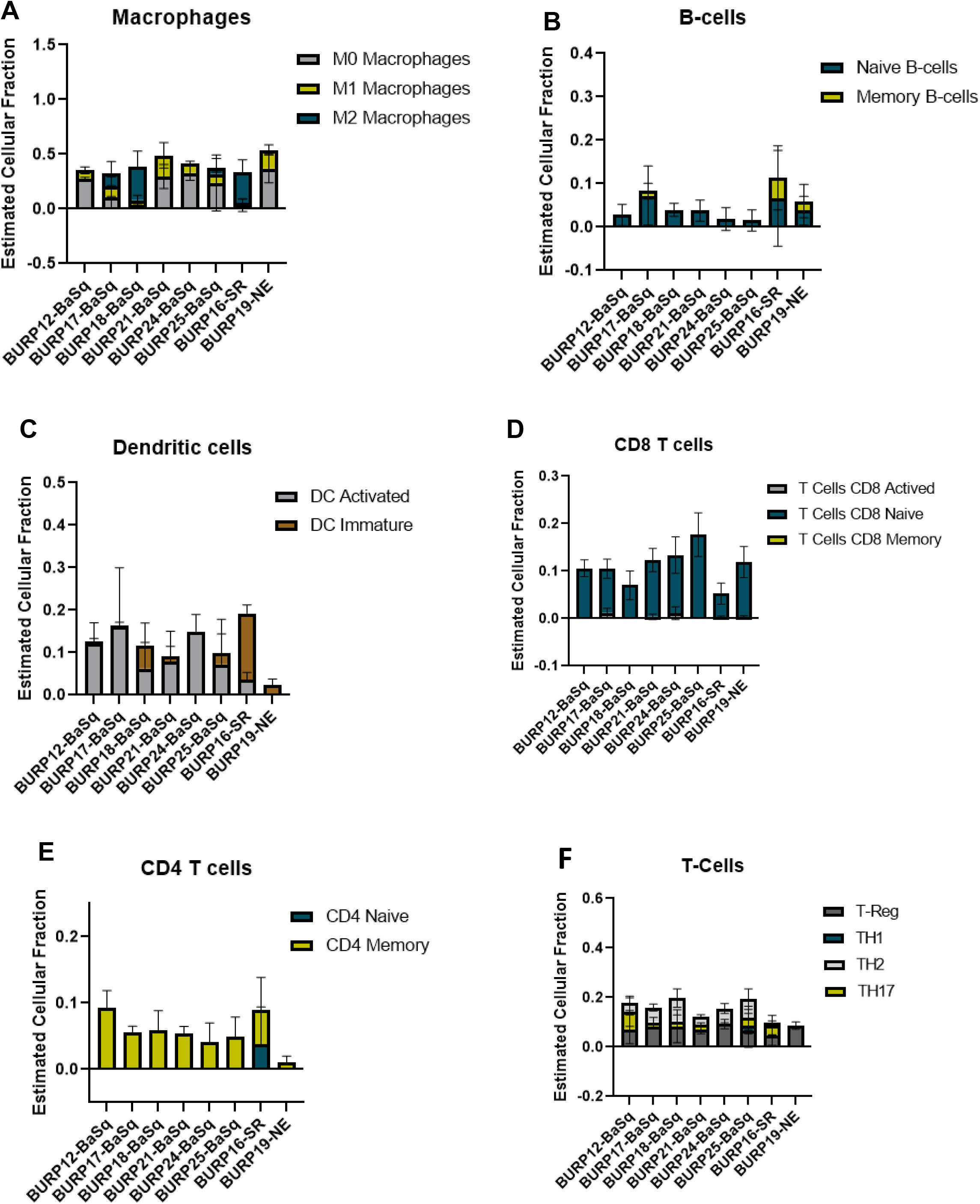
Comparison of theestimated fraction of different immune populations in different BURP tumor lines. **(A)** polarized forms of macrophages. **(B)** naïve and memory B-cells. **(C)** DC immature and activated. **(D)** naïve, activated, memory CD8Tcells **(E)** naive and memory CD4 T-cells. **(F)** T-regulatory cells.

Because improved survival outcomes for bladder cancer are determined by the extent of T cell infiltrates (24, 25), CIBERSORT was used to estimate the fraction of T cells in the BURP lines. The CD8^+^ T cell fraction was overwhelming naïve rather than activated or memory in all three BURP line classifications (Figure 7D). The CD4^+^ T cell fraction was memory in all BURP line classifications except for BURP-16SR, which was a mix of naïve and memory (Figure 7E). Intriguingly, the NE classification (BURP-19NE) had the lowest fraction of CD4^+^ T cells out of all the BURP lines. In the Ba/Sq BURP lines, the fraction of differentiated CD4^+^ T cell helper subpopulations (Th1, Th2, Th17, Treg) was consistently made up of Treg and Th2 with typically fewer Th17, and extremely few if any Th1 cells (Figure 7F). BURP -16SR had Th17 and Treg fractions. In stark contrast, BURP-19NE had almost exclusively a Treg fraction.

Overall, the immune contexture for the four Ba/Sq BURP lines is a mix of pro- and anti-tumorigenic phenotypes. The immune composition in the BURP-16SR tumor line trends towards pro-tumorigenic with a memory B cell signature, and in the BURP-19NE line is pro-tumorigenic with immune exclusion. The immune contexture of each molecular subtype of the BURP lines shows some distinctive features observed in different molecular subtypes of human muscle-invasive bladder cancer. Hence, the BURP lines represent unique models to study the impact of molecular subtype and sex on therapy outcomes including those with an immune component.

### Cisplatin response of Ba/Sq and Stroma-Rich BURP tumors

Since tumors with a stroma-rich molecular subtype have higher survival when treated with neoadjuvant therapy (1), we compared tumor growth of the stroma-rich BURP-16SR to the basal/squam line BURP-24BaSq when treated with cisplatin (Figure 8 A,B). Mice were treated with 10mg/kg of cisplatin and I.V. once a week for four cycles of treatments. Tumor growth was monitored by caliper measurements during treatment (Figure 8 A,B). The BURP-16SR line responded to cisplatin treatment with a decrease in tumor size from the second dose which was sustained throughout the treatment period. In contrast, the BURP-24BaSq line had no difference in the tumor growth compared to vehicle control. BURP-16SR (cisplatin responder) and BURP-24BaSq (cisplatin non-responder) also differ in their immune phenotype (Figure 7 A,C,E,F). BURP-16SR has a high fraction of polarized M2 macrophages, immature DCs, and a mix of naïve and memory CD4^+^ T-cells with a low Th2 differentiated CD4 T-cell fraction. In contrast, BURP-24BaSq had a high fraction of non-polarized M0 macrophages, activated DCs, and Treg and Th2.

**Figure 8:**
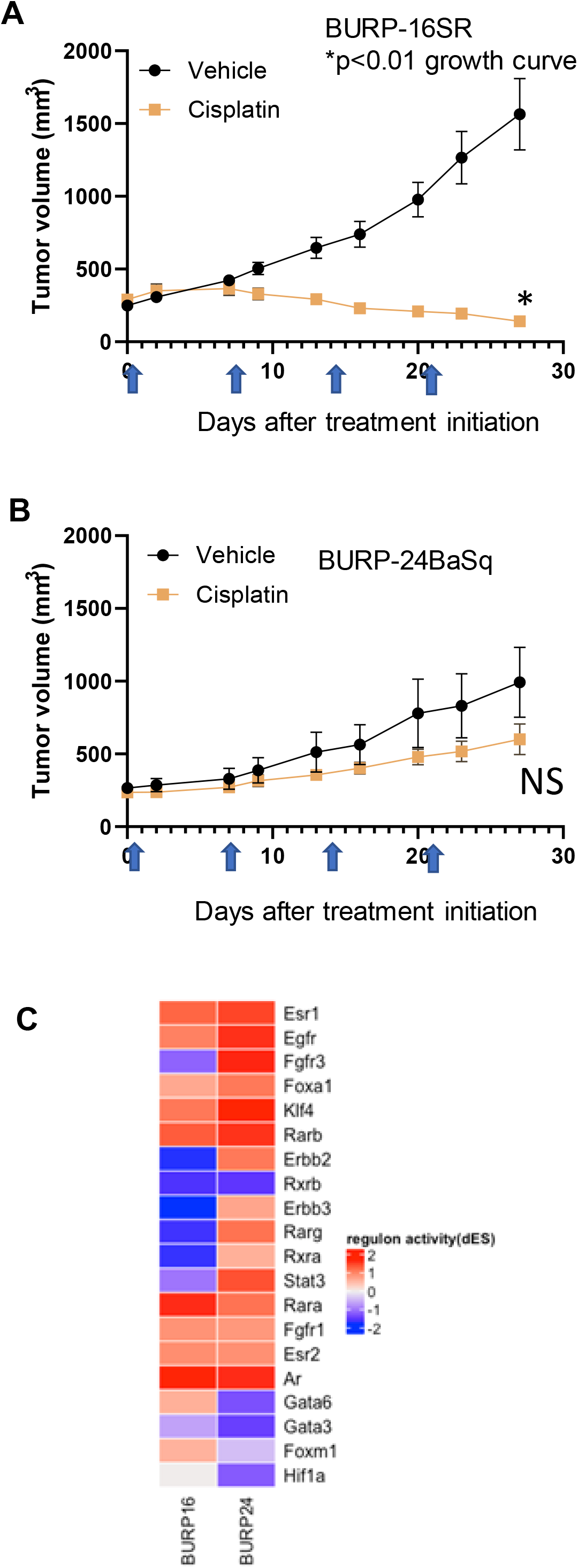
Response of BURP-16SR and BURP-24BaSq tumor lines to cisplatin treatment and regulon activity. **(A) and (B)** Cisplatin treatment (10mg/kg, IV) was initiated when tumors reached 200 mm^3^. Mice were treated on days 0, 7, 14, and 21 (indicated by blue arrows) and tumor growth is determined by caliper measurements twice a week. BURP-16SR n=10 each cohort, * growth curve p<0.01 unpaired t-test. B. BURP-24BaSq n=9 cisplatin and n=8 vehicle cohorts, NS= not significant difference in growth curve p=0.10 unpaired t-test. **(C)** Unsupervised clustering of regulon activity of BURP-16SR and BURP-24BaSq models. The regulons evaluated are identified to be relevant to bladder cancer as represented in Kamoun et al. The color represents differential regulon activity, with red representing active status and blue representing inactive status.

The differential regulon activity was assessed in the two BURP lines to elucidate the molecular underpinnings of the differences between the BURP-16SR and BURP-24BaSq tumor lines (Figure 8 C). The regulon activity represents a network of differential gene activity for the master regulators identified for bladder cancer by Kamoun et al. (1). Interestingly, the cisplatin non-responsive BURP-24BaSq has high regulon activity of Fgfr3, Egfr, and Erbb2. In support of this finding, these three regulons control kinase survival pathways, which are known to confer resistance to cisplatin treatment (26, 27). The histological, molecular, immunological and differences in treatment outcome between the different tumor lines make the BURP models valuable tools in evaluating different targeted therapeutics including those with an immune component for bladder cancer patients across different molecular subtypes.

## Discussion

Eight tumor lines were established from BBN carcinogen exposed mice in an immune competent C57BL/6 mouse strain background. BBN induced tumors in mice closely resemble the clinical disease in terms of molecular subtype and immune contexture. Previous reports by Van Batavia et al (28), indicated that based on the cellular origin of the primary tumor, BBN can induce histologically variant tumors. The BURP lines have different histological features, molecular classifications, and composition of immune infiltrates all of which reflects the variability observed in the BBN-treated tumors.

The BURP lines cluster into three main classifications by transcriptomic evaluation: Basal/Squam, stroma-rich and neuroendocrine. None of the BURP tumor lines represent the luminal subtypes. The six tumor lines of the Ba/Sq subtype (Cluster 1) include 3 male and 3 female lines and will be a valuable tool in evaluating gender differences in treatment response and effect within the basal/squam molecular classification. Although the 6 tumor lines were all classified as Ba/Sq, there was variable expression of the differentiation markers between the six lines. BURP-12, -21, -24 and -25 were positive for TP63 expression, whereas BURP-17 and -24 were negative for cellular TP63 expression. Clinically, TP63 is closely associated with poor outcomes in basal subtype, but is associated with improved survival in the luminal subtype (29). Interestingly, BURP-17 and BURP-24 were positive for FOXA1 and GATA3, which are usually indicators of luminal differentiation when co-expressed with KRT20 (18). Thus, the variability within the Ba/Sq subtype provides a wide array of models within a specific molecular subtype. BURP-16SR and BURP-19NE are two lines that represent stroma-rich and neuroendocrine molecular subtypes. Clinically, patients with a tumor having a stroma- rich classification had a relatively better prognosis (Median OS = 3.8 years) compared to patients with a neuroendocrine classification (median OS = 1 year) (1). Little is known about the biology of these relatively rare molecular subtypes in bladder cancer and thus these models can be used to evaluate not only the treatment differences between the molecular classifications but can also be used for identifying novel targets of treatments for the different subtypes.

The molecular subtype of human bladder cancer associates with a distinct immune contexture that predicts outcome (1). Evaluation of the immune contexture of the BURP tumor lines by applying the machine learning algorithm CIBERSORT to transcriptomic data revealed that the immune profiles of molecular subtypes have some features that recapitulate the human bladder cancer subtypes. The BURP lines that sort into the Ba/Sq classification are a mix of expressing immune markers associated with an anti-tumor or a pro-tumorigenic phenotype. Despite being heavily infiltrated with CD8^+^ T cells, human Ba/Sq tumors do not respond to immunotherapy as well as less heavily infiltrated tumors (1). M2 macrophages are thought to be responsible for the poor response to immunotherapy (30). BURP-17, -18, and -25 have a relatively high fraction of M2 macrophages and could be representative models to evaluate the role of M2 macrophages in treatment resistance. These models can also be used to test novel therapeutic approaches for the treatment of muscle-invasive bladder cancer. The stromal rich BURP-16 line had the highest B cell signature, which included both naïve and memory B cells, among all the tumor line classifications and agrees with the clinical observation that the stromal rich subtype was marked by B cell lineage expression signature (1). For the neuroendocrine classification, both human bladder tumors and BURP-19NE had limited expression of immune markers, which supports the notion that these tumors are immune excluded. Although NE tumors have the poorest outcome to chemotherapy, a recent study intriguingly showed an extraordinarily high response to immunotherapy (14). For all the BURP lines, the macrophage signature was the highest among all the immune cell signatures and could be due to the fact that macrophages are the most populous immune cell within the bladder (30). Also of note is that although the BURP tumor lines were derived from immunocompetent hosts, they had little to no presence of an activated or memory CD8^+^ T cell signature which could be the result of selecting serial passaged clones that escape T cell immunity. Although the BURP tumor lines do not capture the full range of the immune profiles observed in the clinical disease, these lines represent a wide range of models to study different treatment regimens for distinct immune contextures, as well as models to evaluate the relationship between tumor immune infiltrates and the various molecular subtypes.

Cisplatin is the current standard of care in the neoadjuvant setting. Based on retrospective clinical observations, the different molecular subtypes are prognostic of response (1, 7, 14, 31). Therefore, the cisplatin response was evaluated in two BURP lines with molecular classifications whose response to cisplatin differs in the clinical setting. Tumor growth and final tumor size was inhibited by cisplatin in BURP-16SR. In contrast, BURP-24BaSq was resistant to cisplatin treatment. The cisplatin response in the BURP lines is in agreement with the clinical observation, where patients with stroma-rich subtype tumors have better prognosis as compared to basal/squam subtype (1). To dive further into evaluating the molecular differences between BURP-16SR and BURP-24BaSq, regulon analysis was performed on the transcriptomic data. The regulon differential analysis showed high activity of regulons known to confer resistance to cisplatin therapy (Egfr, Erbb2 and Fgfr3) in the cisplatin non-responsive BURP-24BaSq. This data provides important mechanistic insight into the cisplatin resistance of the BURP-24BaSq tumor line.

Taken together, the BURP tumors can be used to evaluate the molecular differences driving therapeutic response and hence may be important tools in identifying novel targets or combination regimens to improve the clinical response in particular molecular subtypes. Additionally, since the BURP lines grown in syngeneic, immunologically intact hosts, these lines offer an immune intact system to study ways of switching the immune microenvironment from tumor supporting to tumoricidal as a means to increase the anti-cancer therapeutic response. Hence, the eight different tumor lines are valuable tools to study the biology of different molecular subtypes and tumor-immune dynamics of bladder cancer in both males and females for the purpose of identifying novel targets and treatment combinations.

## List of Abbreviations

BaSq: basal/squamous
BBN: N-butyl-N-(4-hydroxbutyl nitrosamine)
BSA: Bovine serum albumin
BURP: BBN-induced Urothelium Roswell Park
EMT: Epithelial mesenchyme transition
ETM: Experimental tumor model
DCs: Dendritic cells
GSR: Genomic shared resource
IACUC: Institutional Animal Care and Use Committee
IHC: Immunohistochemistry
LASR: Laboratory Animal Shared Resource
LumNS: Luminal Non-Specified
LumP: Luminal Papillary
LumU: Luminal Unstable
MIBC: Muscle-invasive bladder cancer
MDS: Multidimensional scaling
NE: neuroendocrine
NMIBC: Non-muscle invasive bladder cancer
PDX: Patient-derived xenograft
RIN: RNA integrity number
SR: Stromal-rich
TILs: tumor-infiltrating lymphocytes
Tregs: T regulatory cells

## Acknowledgements

The authors thank Kevin Eng for thoughtful discussions, Jordon McDonald with help formatting figures and Roswell Park shared resources: Genomic shared resource (GSR) for genomic analysis, Experimental tumor model (ETM) for animal *in vivo* work, LASR for animal housing and veterinary oversight.

## Declarations

Ethics approval and consent to participate: No human specimens were used in the studies. All animal experiments were conducted and approved under our Institutional Animal Care and Use Committee (IACUC) protocol at Roswell Park.

